# Evidence for rapid hydrolysis of shoot-derived sucrose using an ultrasensitive ratiometric Matryoshka-type MGlucoMeter sensor

**DOI:** 10.1101/2025.09.27.678933

**Authors:** Yuuma Ishikawa, Nora Zöllner, Susanne Paradies, Wolf B. Frommer

## Abstract

To enable sensitive *in vivo* monitoring of the glucose transport and metabolism, we developed a series of ultrasensitive and ratiometric genetically encoded nanosensors by inserting a Matryoshka dual fluorophore cassette consisting of cpsfGFP and LSSmApple into the glucose binding protein ttGBP from *Thermus thermophilus*. The initial MGlucoMeter1.0 was subjected to an alanine scan of the hinge region producing more sensitive MGlucoMeter2.6 with a glucose-induced ΔF/F_0_ change of 3.0, an affinity for glucose of 15 µM, and an approximate detection range of 1.1-216 µM. To generate variants suitable for *in vivo* measurements, a series of affinity mutants was generated by mutating two histidines predicted to be involved in substrate binding. MGlucoMeter2.6-353n, MGlucoMeter2.6-15µ, MGlucoMeter2.6-700µ, MGlucoMeter2.6-1m, and MGlucoMeter2.6-7m cover a detection range between ∼40 nM - 55 mM. When expressed from a ubiquitous promoter in the cytosol of the Arabidopsis gene silencing mutant *rdr6*, MGlucoMeter2.6-1m reports time- and concentration-dependent accumulation of glucose in seedling roots. The sensor also detects rapid hydrolysis of shoot-derived sucrose in the root tip.

## Introduction

While metabolomics provides quantitative information on the steady state levels of metabolites, it lacks spatial and time resolution. An alternative is the visualization of metabolite dynamics with subcellular resolution using genetically encoded nanosensors. The first sugar sensors were developed by sandwiching a sugar binding proteins from the family of bacterial periplasmic binding proteins between two variants of the green fluorescent proteins that had features allowing for Förster Resonance Energy Transfer (Deuschle *et al*., 2005; Fehr *et al*., 2003; Fehr *et al*., 2002; Lager *et al*., 2006). These sensors were successfully used for monitoring glucose dynamics in bacterial and yeast cells, mammalian cells and in the plant Arabidopsis (Bermejo, Haerizadeh, *et al*., 2011; Chaudhuri *et al*., 2011; Chaudhuri *et al*., 2008; Deuschle *et al*., 2006; Kaper *et al*., 2008). The glucose sensors were also successfully deployed for genetic screens to identify networks involved in the regulation of glucose transport and for studying the transfer of glucose across ER membranes (Bermejo *et al*., 2013; Bermejo, Ewald, *et al*., 2011; Bermejo *et al*., 2010; Fehr *et al*., 2005; Takanaga *et al*., 2008). While the FRET-based approach yielded sensors suitable for *in vivo* measurements, the sensitivity of the sensors was limited due to the intrinsic coupling of the relative change in the emission of the two fluorophores. Roger Tsien had developed an alternative approach using a circularly permutated GFP as an intensiometric reporter element (Baird *et al*., 1999). Although the sensitivity of the intensiometric cpGFP-based sensors was higher, the FRET sensors had one major advantage: they were ratiometric and thus not impacted by changes in the sensor levels. To overcome the limitation, calcium and ammonium transporter activity sensors were generated that contained a nested large Stokes shift fluorophore (LSSmOrange) as a reference, enabling ratiometric measurements (Ast *et al*., 2015; Ast *et al*., 2017). The calcium sensor was further improved by replacing LSSmOrange with the better suited LSSmApple (Ejike *et al*., 2024). Since proteins from thermophilic organisms are more robust, the glucose-binding protein from *T. thermophilus* had previously been used to generate an intensiometric glucose sensors by inserting cpGFP into ttGPB to produce iGlucoSnFR (Keller *et al*., 2021).

Here, a series of ultrasensitive glucose sensors (MGlucoMeter) was generated using the Matryoshka technology. For this purpose, the previously developed Matryoshka cassette, consisting of the superfolder variant cpsfGFP as a sensory domain and LSSmApple as a reference fluorophore (Ejike *et al*., 2024; Pedelacq *et al*., 2006), was inserted into the glucose binding protein ttGBP from *Thermus thermophilus*. The initial sensor was further optimized by linker mutagenesis, and site directed mutagenesis was used to produce a series of affinity mutants. MGlucoMeter2.6-1m with an affinity of 1 mM for glucose was then used to monitor the accumulation of glucose in Arabidopsis roots as well as the accumulation and transport of glucose in Arabidopsis seedling roots after supply of sucrose to shoots.

## Material and Methods

### Construction of MGlucoMeter

MGlucoMeter2.0 were generated from the initial construct MGlucoMeter1.0 (www.molecular-physiology.hhu.de/en/resources-1/mglucometer-10), which carries an insertion of the cpsfGFP-LSSmApple cassette from GA-MatryoshCaMP6s into the hinge region of ttGBP from *Thermus thermophilus* in the pRSETb T7 expression vector (Thermo Scientific; V35120) (Ejike *et al*., 2024; Cuneo *et al*., 2009) (Fig.1a, Table1). The expressed gene product (P0328) of ttGBP lacks the 21-bp leader sequence (Cuneo *et al*., 2009). An alanine scan of the linker and hinge region (amino acids targeted left linker/hinge region: DSDPSKYPASH; right linker/hinge region FNNPNAYGQSAM) was performed by inverse PCR (Fig. 1a, Table 1). PCR products were amplified using PrimeSTAR GXL (Takara Bio; R050B) (Table S1). The resulting product was cleaned up by gel extraction using the NucleoSpin column kit (Macherey-Nagel; 740609). Affinity variants were generated by introducing single point mutations into ttGBP. All sequences were confirmed by DNA sequencing (Microsynth SEQLAB). Plasmids containing the MGlucometer2.6 affinity mutants are available from AddGene (www.addgene.org).

**Table 1.**
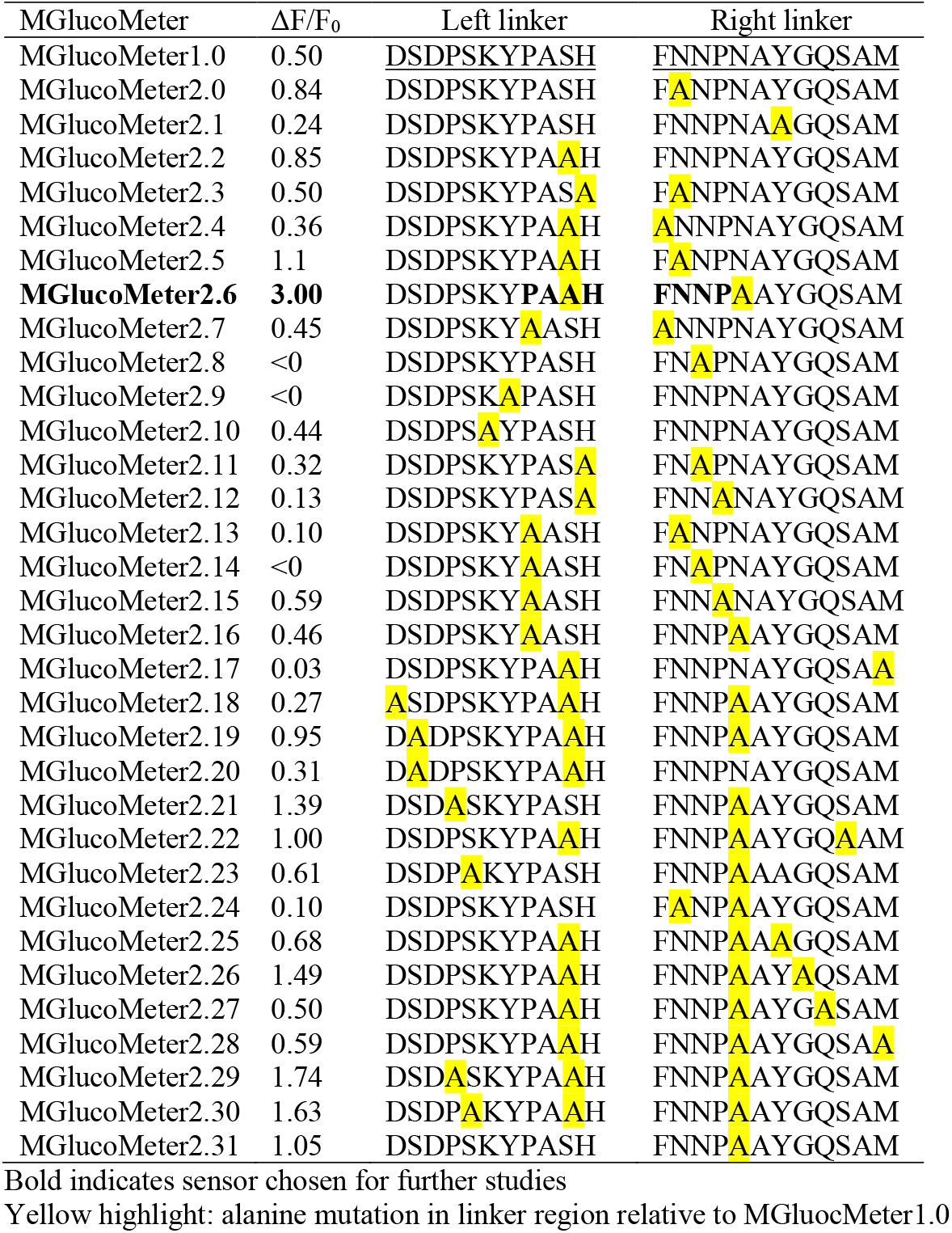
Sequence of linker/hinge sequence subjected to alanine scanning of MGlucometer variants.

**Figure 1:**
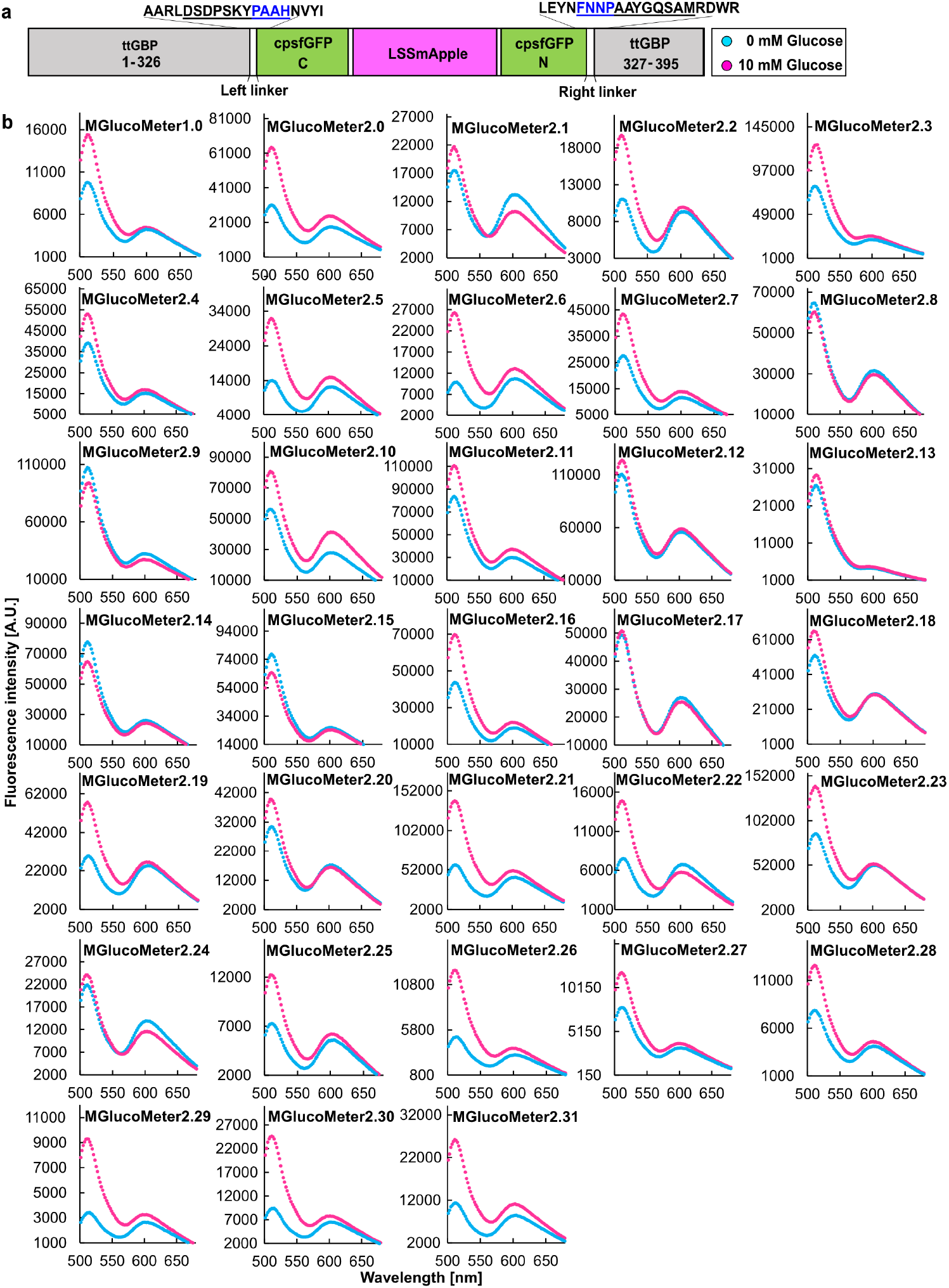
*In vitro* screening of MGlucoMeter variants with increased sensitivity. **a**. Construct map of MGlucoMeter2.6 carrying an insertion of the Green-Apple cassette (GA; cpsfGFP-LSSmApple) in ttGBP. The amino acid residues substituted for MGlucoMeter are indicated as underlined. Four amino acid linker sequences connecting the Green-Apple cassette and ttGBP are indicated with blue underline. **b**. The fluorescence responses of 32 purified MGlucometer variants, generated by the alanine scan, were tested *in vitro* for changes upon addition of 10 mM glucose.

### Expression and purification of the MGlucoMeter

*E. coli* BL21 (DE3) [*fhuA2 [lon] ompT gal (λ DE3) [dcm] ΔhsdS; λ DE3 = λ sBamHIo ΔEcoRI-B int::(lacI::PlacUV5::T7 gene1) i21 Δnin5*] (New England Biolabs; C2527I) was transformed with pRSETb containing MGlucoMeter plasmids. Single colonies were inoculated into 3 mL of LB medium (Duchefa Biochemie; 4800-94-6) containing 50 μg/mL carbenicillin (Sigma-Aldrich; C1389) in glass tubes (Hardy diagnostics; 1517) sealed with φ15/16 mm silver aluminum caps (Lüdi; LUDI184010631) and incubated at 37 °C with shaking at 200 rpm for 14-18 h. 2 mL of the starter culture were transferred into 100 ml LB medium containing 50 μg/mL carbenicillin disodium, 0.2% D-lactose (Sigma-Aldrich; 64044-51-5), and 0.05% D-glucose monohydrate (Thermo Scientific Chemicals; 14431-43-7) in 500 mL Erlenmeyer flasks without baffles (SciLahware; 1135/26D) sealed with φ 37/39 mm silver screw caps (Roth selection; K395.1). After 2 h of incubation at 37 °C with shaking at 200 rpm, cultures were transferred to 20 °C incubator with shaking at 200 rpm and incubated for 48 h. Bacterial cultures were cooled on ice and centrifuged at 3,790 × g for 10 min at 4 °C. The cell pellet was stored at −20 °C for at least 14 h. After thawing on ice, the pellet was resuspended in 1.5 mL of 20 mM MOPS (Roth; 1132-61-2), pH 7.0. To prevent sensor degradation, one cOmpleteTM ULTRA Tablet Mini protease inhibitor cocktail (Merck; 1183670001) was added per 100 mL of all purification buffers. Cells were lysed by sonication at 50 amplitude (QSonica Q700 Sonicator), with a total processing time of 45 s (3 s pulse on, 8 s pulse off). The sonicated sample was centrifuged at 15,871 × g for 10 min at 4 °C to remove cellular debris. His-tagged MGlucoMeter sensor proteins were purified using Ni-NTA agarose beads (Qiagen; 30210). Prior to lysate application, 1.5 mL of Ni-NTA agarose beads was loaded onto a column and washed with 10 mL of 20 mM MOPS, pH 7.0. After the loading of the lysate, the column was washed again with 10 mL of 20 mM MOPS, pH7.0. His-tagged MGlucometer protein was eluted with 1.5 mL of 250 mM imidazole (Sigma-Aldrich; 1467-16-9) and 20 mM MOPS, pH 7.0. Protein concentration was determined by a NanoDrop One (Thermo Scientific). Purified sensors were incubated at 4 °C for at least 14 h to ensure maturation of the fluorescent protein.

### Fluorimetric analysis of MGlucoMeter2.0 sensors

For each well, 10 μL of sensor solution was mixed with 190 μL of buffer. Ligand titrations were performed using a TECAN Spark microplate reader (Tecan Austria GmbH; 704004367). Steady-state fluorescence spectra were recorded with excitation wavelength at 453 ± 20 nm and emission from 499 to 700 nm in 2 nm steps, using a 50% mirror. Measurements were recorded in the top-reading mode with a gain of 80–130 and the temperature controlled at 27 ± 0.5 °C. Spectra were background-subtracted using 20 mM MOPS containing 250 mM buffer (pH 7.0). Analysis was performed in 96-well clear, flat-bottom, microplate (Corning; 3370). Fluorescence emission ratios were calculated as cpsfGFP emission intensity (λ_ex_ 480 nm, λ_em_ 515 nm) divided by LSSmApple emission intensity (λ_ex_ 460 nm, λ_em_ 600 nm). The baseline ratio defined as *R*_0_ and the ratio at increasing ligand concentrations as R. Data were normalized accordingly;

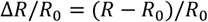

Data analysis and visualization of the data were performed using Excel, Origin Lab 2020, and Affinity Designer. Sigmoid curve fitting was performed with MyCurveFit (https://mycurvefit.com). Values are reported as the mean ± standard error of Δ*R*/*R*_0_ or Δ*F*/*F*_0_ from n = 3 biological replicates (*n* = 3 technical replicates per biological replicates) out of two independent titrations. The dissociation constant (*K*_d_) was defined as the ligand concentration at 50% Δ*R*_*max*_/*R*_0_ in the fitted sigmoid function.

### Cloning of MGlucometer2.6 constructs for expression in plants

MGlucometer2.6-1m was subcloned into pAY367-HTv443-UBQ10-Hspt binary vector (provided by Dr. A. Yoshinari, Institute for Transformative Biomolecules, ITbM, Nagoya University, Nagoya, Japan), which contains the Arabidopsis UBQ10 promoter and Hsp18.2 terminator. To insert MGlucometer2.6-1m, pAY367-HTv443-UBQ10-Hspt was digested with *AscI* and *ApaI* restriction enzymes. Ligation reaction was carried out in the solution mixture containing 40 fmol DNA of the entry clones, 1x T4 DNA ligase buffer (Thermo Fisher; B69) and 5U T4 DNA ligase (Thermo Fisher; EL0012) in a total volume of 20 µL (Thermocycler: 22 ℃ for 10 minutes). The Arabidopsis gene-silencing mutant *rdr6-11* (Peragine *et al*., 2004; Deuschle *et al*., 2006) was transformed using the floral dip method. Transformed plants were selected on 50 µg/ml kanamycin and 2.35 mM MES containing ½ salt strength Murashige and Skoog (MS) medium (Duchefa Biochemie; M0255), adjusted to pH 5.7 with KOH, and solidified with 1% agar (Merck; 05040).

### Plant growth conditions

Seeds were surface sterilized with 75% ethanol containing 0.08% Tween20, sown on ½ salt strength MS medium containing 2.35 mM MES, adjusted to pH 5.7 with KOH, and solidified with 1% agar. Plants were sealed with micropore tape and vernalized at 4 ℃ for 1 day. Plants were grown at 22 ℃ for 5 days after germination under the long day conditions (16 hours light/8 hours dark, light intensity was 110 µmol m^-2^ s^-1^).

### Root uptake kinetics measured with an automated perfusion system

MGlucometer2.6 seedlings were germinated, and 10-day-old seedlings were transferred to ½ salt strength MS medium in a 60 µ-Dish (Ibidi; 81158) and immobilized using double side adhesive tape (Tesa Tape; 05338). Seedlings were acclimated for 20 min by perfusion with mock buffer (¼ salt strength MS containing 2.35 mM MES, pH 5.7) in the recording room. A peristaltic pump was used (Fisher Scientific; Cytiva 18-1110-91, flow rate; 2 ml/min) to perfuse the filtered ¼ salt strength MS containing 2.35 mM MES (pH 5.7) liquid media to the whole seedling. Fully automated buffer exchange to 1 mM glucose containing ¼ salt strength MS containing 2.35 mM MES (pH 5.7) liquid media was achieved using valve controller (Automate scientific; ValveBank Controllers). The holdup time (delayed arrival of buffer in a fluid reservoir) was assessed in the perfusion set-up using a fluorescent dye (ATTO 390; ATTO-TEC GmbH, AD 390-25). Holdup time was calculated by taking the saturated fluorescence intensity as 100% and determining the time that required to reach 85% or 15% of that fluorescence intensity. An Olympus IXplore SpinSR spinning disk confocal microscope was used for imaging. Samples were excited with a 488 nm laser (Obis; 100mW, used at 10-30% power). The microscope was equipped with a Yokogawa confocal scanning unit (CSU)-W1 SoRa micro-lensed pinhole disk and an Olympus UAPON 10× UPL SAPO10×2. Emission intensities were collected in sequential stacks using a 525/50nm emission filter for GFP, 617/73 nm emission filter for LSSmApple. For detection, an Andor iXon Ultra 888 electron-multiplying charge coupled device (EMCCD) was used. Z-stacks were set by using a Mad City Labs Z-axis piezo nano positioner with 300 nm travel range (OLY-S1023-Nano-ZL300-OSSU). Full Z-stack acquisitions for each channel were performed at a frame rate of 1 min. The exposure time for each channel was 50 msec, with 4×4-pixel binning.

### Quantitative imaging of glucose in roots in response to sucrose addition to shoots

Seedlings were acclimated for at least 2 h on 1% agar plates containing ½ salt strength MS and 2.35 mM MES (pH 5.7), sealed with micropore tape, in the recording room. Seedlings were placed on 1% agar plates containing ½ salt strength MS and 2.35 mM MES (pH 5.7) in a glass bottom chamber (Ibidi; µ-slide 1 well glass bottom). A flat PCR tube cap filled with 2M sucrose containing ¼ salt strength MS containing 2.35 mM MES (pH 5.7) liquid media was placed in the glass bottom chamber, with the tip of cotyledon submerged in the sucrose solution (Fig. S2). Quantitative glucose imaging was performed using a Zeiss AxioZoom V16 zoom microscope equipped with a X-Cite XYLIS LED Illumination System (XT720L), a 1 × objective lens (PlanNeoFluar Z 1 × /0.25 NA, FWD 56 mm, Zeiss, Oberkochen, Germany), and a Hamamatsu ORCA Flash4.0 CMOS camera. Zoom magnification was set between 7× and 10×, and pixel binning was set to 2 × 2 or 4 × 4-pixel binning for acquisitions. cpsfGFP fluorescence was detected using a filter cube with a 488/10 nm excitation filter, a 491 nm long-pass dichroic mirror, and a 519/26 nm emission filter. LSSmApple fluorescence was detected using a filter cube with a 488/10 nm excitation filter, a 514 nm long-pass dichroic mirror, and a 605/50 nm emission filter. A motorized *xy* stage (Zeiss) was used for time-lapse tiling acquisitions. Tiles were stitched using ZEN Blue 2.6 software (Zeiss). *Z*-step sizes were set equal to the optical section thickness (2-4 µm).

### Image analyses

Segmentation and labelling were performed with the FRETENATOR plugins in Fiji (Schindelin *et al*., 2012). Segmentation settings were optimized for each experiment. The GFP channel was used for segmentation. Watershed algorithm was used for the image segmentation. Difference of Gaussian kernel size was determined empirically due to different magnifications, resolutions and amount of noise. As a default, Otsu thresholds were used for segmentation in FRETENATOR (Rowe *et al*., 2022).

## Results and Discussion

### Development of new ultra-sensitive Matryoshka-type sugar sensors

Proteins from thermophiles often exhibit enhanced stability. For generating a Matryoshka-type glucose sensor, the glucose binding protein from *Thermus thermophilus*, for which crystal structure information is available, was selected as a recognition element (Cuneo *et al*., 2009; Keller *et al*., 2021). The glucose binding protein ttGBP had been used to develop the intensiometric glucose sensor GluSnFR by inserting the *T. thermophilus* GBP lacking the leader peptide and a linker combination left linker-ProAla/right linker-AsnPro (L1-PA/L2-NP) between residues 326 and 327 of ttGBP (Keller *et al*., 2021, Supplementary Text). Analogous to the construction of GluSnFR, but here with the Matryoshka cassette consisting of cpsfGFP carrying an insertion of the Large Stokes Shift (LSS) mApple (LSSmApple) that had been used to generate MatryoshCaMP6s (Ejike *et al*., 2024) was inserted into ttGBP without the N-terminal signal sequence to generate MGlucoMeter1.0 (https://www.molecular-physiology.hhu.de/en/resources-1/mglucometer-10). The linkers that flank the fusion site between cpsfGFP and the recognition element are thought to be important for ESPT^22^ (De Michele *et al*., 2013). To further increase the sensitivity and detection range for *in vivo* measurements, the codons for amino acid pairs in the linker sequence flanking the Matryoshka cassette in MGlucoMeter1.0 were mutated. An alanine scan was performed corresponding to 19 amino acids surrounding the cpsfGFP insertion site; i.e. the hinge region of the glucose-binding domain, and linker sequences (FNNPNAYGQSAM/DSDPSKYPASH); (Fig. 1a, Table 1). A set of 32 mutants carrying single, double or triple alanine replacements were characterized (Table 1). Fluorometric analysis of the glucose-induced ratio changes of the series indicated that some of the sensors lost responses, some showed altered affinities (Fig. 1b). We selected one variant that showed the highest sensitivity MGlucoMeter2.6 (ΔF/F_0_ 3.0) for generating a series of affinity variants.

### Generation of an affinity variants for MGlucoMeter2.6

Previous studies had shown that FLIPglu600µ with an affinity for glucose of 600 µM was well suited for recording glucose responses in Arabidopsis roots, we generated a series of affinity mutants of MGlucoMeter2.6 by site directed mutagenesis of candidate residues that interact with the glucose molecule: H66 and H348 (numbering based on the previously published sequence of ttGBP (Fig. S1; Supplementary Text)(Chaudhuri *et al*., 2008; Cuneo *et al*., 2009; Keller *et al*., 2021). The resulting mutants exhibited affinities (K_d_) for glucose of approximately 355 nM, 717 µM, 1 mM, 7.3 mM, respectively (Fig. 2, 3 and Table 2). All affinity variants showed higher sensitivity (ΔF/F_0_ 1.90-3.69) compared to the FLIPglu (Fig. 3, Table 3). Notably, the detection range of the MGlucometer2.6 sensors is substantially larger compared to FLIPglu, which covered less than two orders of magnitude (Table 3) (FLIPglu-600µ: 65.4-5301µM)(Fehr *et al*., 2003). Analysis of the substrate specificity showed that glucose had the highest affinity followed by galactose, whereas other sugars showed much lower affinities (Fig. 4, Table 4). Of note, MGlucoMeter2.6-1m used below for *in planta* analyses, did not show detectable *in vitro* responses to the addition of sucrose (Fig. 4).

**Table 2.** Affinity and mutation of MGlucoMeter2.6 variants.

| MGlucoseMeter | Mutation | K<sub>d</sub> glucose |
| --- | --- | --- |
| MGlucoseMeter2.6a | H66A + H348R | n.r. |
| **MGlucoseMeter2.6-700μ** | **H66A + H348N** | **717 ± 0.5 μM** |
| MGlucoseMeter2.6b | H66A + H348D | n.r. |
| **MGlucoseMeter2.6-7m** | **H66A + H348C** | **7.3 ± 0.5 mM** |
| MGlucoseMeter2.6c | H66A + H348Q | 42.9 mM |
| MGlucoseMeter2.6d | H66A + H348E | >150mM |
| **MGlucoseMeter2.6-1m** | **H66A + H348A** | **1 ± 0.01 mM** |
| **MGlucoseMeter2.6-353n** | **H66H + H348A** | **355 ± 2.7 nM** |
| MGlucoseMeter2.6e | H66A + H348K | n.r. |
| MGlucoseMeter2.6f | H66A + H348F | n.r. |
| MGlucoseMeter2.6g | H66A + H348P | 210.5 mM |
| MGlucoseMeter2.6h | H66A + H348S | 65.1 mM |
| MGlucoseMeter2.6i | H66A + H348T | 95.3 mM |
| MGlucoseMeter2.6j | H66A + H348W | n.r. |
| MGlucoseMeter2.6k | H66A + H348Y | n.r. |
| **MGlucoseMeter2.6-15μ** | **H66A + H348H** | **15 ± 0.46 μM** |

Bold indicates sensor chosen for further studies
n.r. no response

**Table 3.** Sensitivity and detection range of MGlucoMeter2.6 variants.

| MGlucoseMeter | ΔF/F<sub>0</sub> | Detection range (10 ~ 90% sat) |
| --- | --- | --- |
| MGlucoseMeter2.6-353n | 1.90 ± 0.06 | 0.04 μM ~ 5.1 ± 0.1 μM |
| MGlucoseMeter2.6-15μ | 3.00 ± 0.13 | 1.1 ± 0.005 μM ~ 216 ± 14.4 μM |
| MGlucoseMeter2.6-700μ | 2.63 ± 0.18 | 46.6 ± 1.6 μM ~ 8.6 ± 1.0 mM |
| **MGlucoseMeter2.6-1m** | **3.69 ± 0.14** | **102.7 ± 0.003 μM ~ 10.0 ± 0.76 mM** |
| MGlucoseMeter2.6-7m | 2.21 ± 0.19 | 0.39 ± 0.03 mM ~ 55.9 ± 11.4 mM |

Bold indicates sensor chosen for further studies

**Table 4.** Substrate selectivity of MGlucoMeter2.6 variants.

| K<sub>d</sub> Sugars | MGlucoseMeter2.6 |  |  |  |  |
| --- | --- | --- | --- | --- | --- |
|  | 353n | 15μ | 700μ | 1m | 7m |
| Glucose | 355 ± 2.7 nM | 15 ± 0.4 μM | 717 ± 0.5 μM | **1 ± 0.01 mM** | 7.3 ± 0.5 mM |
| Xylose | 1.83 ± 0.6 mM | 25.6 ± 2.3 mM | n.r. | **n.r.** | n.r. |
| Maltose | 62.3 ± 0.07 μM | 0.97 ± 0.02 mM | n.r. | **n.r.** | n.r. |
| Trehalose | 0.23 ± 0.06 mM | 0.72 ± 0.02 mM | n.r. | **n.r.** | n.r. |
| Sucrose | n.r. | n.r. | n.r. | **n.r.** | n.r. |
| Raffinose | n.r. | n.r. | n.r. | **n.r.** | n.r. |
| Lactose | 11 ± 2.2 μM | n.r. | n.r. | **n.r.** | n.r. |
| Galactose | 2.8 ± 0.3 μM | 38.2 ± 4.5 μM | 2.87 ± 0.2 mM | **14.9 ± 0.7 mM** | 25.5 ± 2.2 mM |

Bold indicates sensor chosen for further studies
n.r. no response

**Figure. 2:**
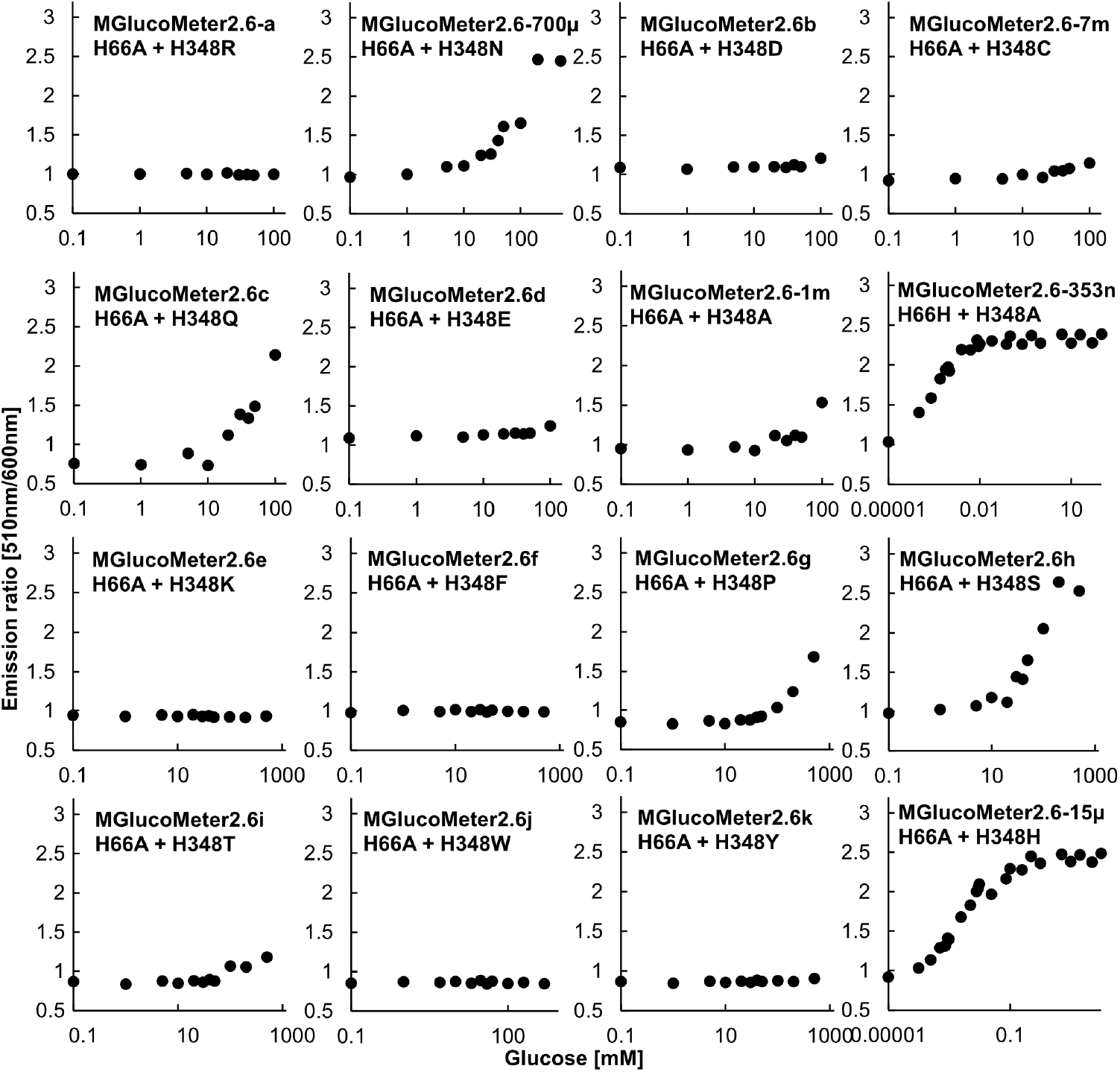
*In vitro* screening of affinity variants based on MGlucoMeter2.6. 16 affinity variants, generated by site directed mutagenesis at H66 and H348, were characterized *in vitro* using the cpsfGFP/LSSmApple emission ratio. Purified protein was titrated with increasing glucose concentrations.

**Figure 3:**
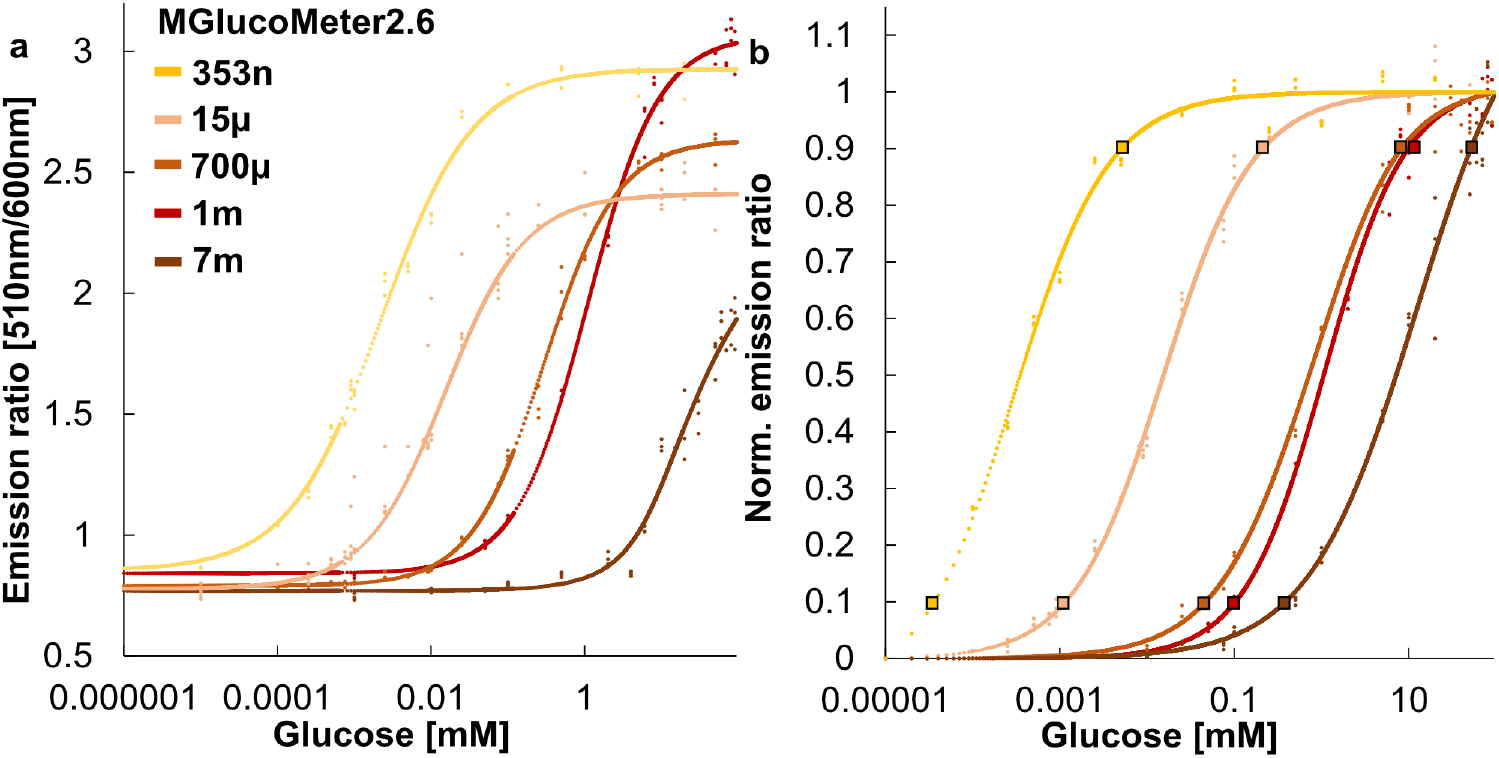
Glucose concentration dependent changes in fluorescence emission ratio of MGlucoMeter2.6 affinity variants. **a**. *In vitro* substrate titrations of MGlucoMeter2.6 affinity variants. MGlucoMeter2.6-353n (H66H + H348A), MGlucoMeter2.6-15µ (H66A + H348H), MGlucoMeter2.6-700µ (H66A + H348N), MGlucoMeter2.6-1m (H66A + H348A), and MGlucoMeter2.6-7m (H66A + H348C) were generated by site directed mutagenesis. The *in vitro* properties of each sensor are summarized in Tables 2 and 3. The mean of three technical replicates is plotted. Experiments were repeated independently two times with comparable results. **b**. Estimation of the detection range of MGlucoMeter2.6 (linear range between 10 and 90% saturation).

**Figure 4:**
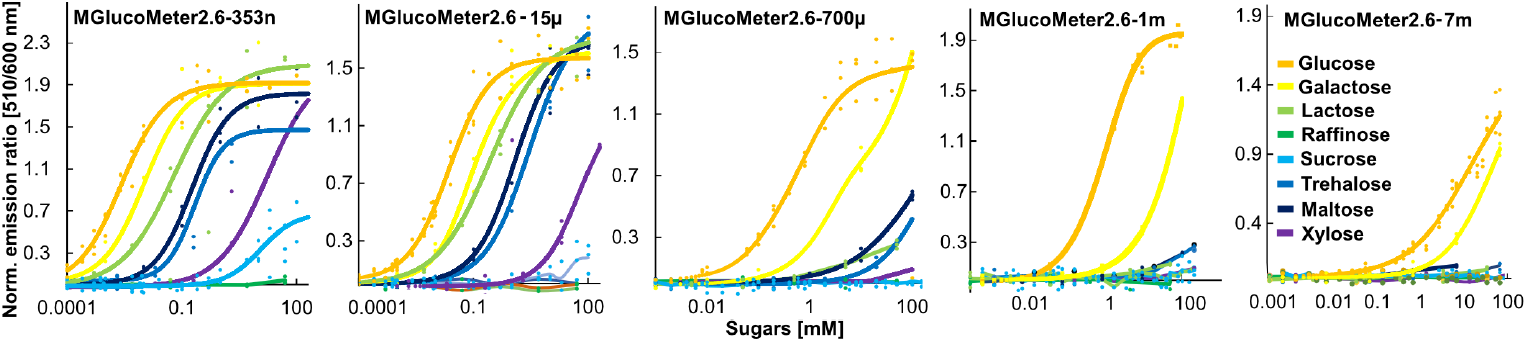
*In vitro* determination of the substrate selectivity of MGlucoMeter2.6 affinity variants. *In vitro* substrate titrations of MGlucoMeter2.6 affinity variants. MGlucoMeter2.6-353n, MGlucoMeter2.6-15µ, MGlucoMeter2.6-700µ, MGlucoMeter2.6-1m, MGlucoMeter2.6-7m were tested to determine the substrate selectivity. The *in vitro* properties of each sensor are summarized in Table 4. The mean of three technical replicates is plotted. Experiments were repeated independently two times with comparable results.

### Monitoring uptake of glucose into Arabidopsis roots

To be able to monitor glucose concentration changes *in planta*, MGlucoMeter2.6-1m was expressed from the *UBQ10* promoter in the cytosol of the Arabidopsis gene silencing mutant *rdr6*. Roots of 10-day-old seedlings mounted in a hand-made perfusion chamber were perfused with square pulses of glucose and the emission change after addition of glucose was monitored under an inverted fluorescence microscope. Within one minute after addition of 1 mM glucose, MGlucoMeter2.6-1m showed a detectable emission ratio change in the root (Fig. 5, Supplementary Video 1). Within about two minutes, the maximal response was reached, which would correspond to reaching the K_d_ of 1 mM, given that previous experiments with FRET glucose sensors had shown that extra- and intracellular levels of glucose rapidly equilibrated (Chaudhuri *et al*., 2008). During further perfusion, the sensor response remained constant, and declined rapidly after replacing glucose-containing buffer with glucose-free buffer, consistent with reversibility of the response.

**Figure 5:**
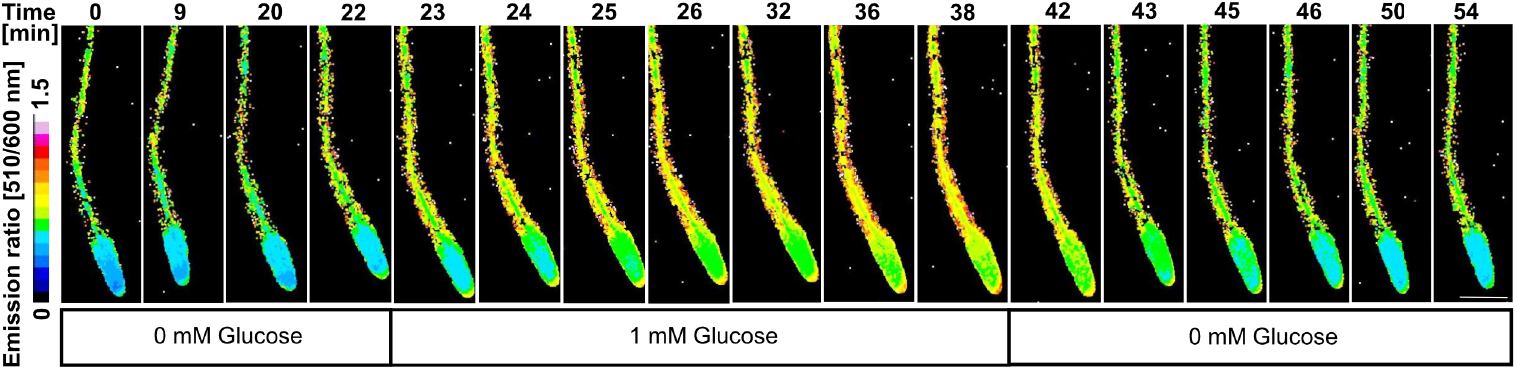
Time-dependent accumulation of glucose in Arabidopsis roots as monitored with MGlucoMeter2.6-1m to perfusion of roots with 1 mM glucose. Time-dependent *in vivo* response of MGlucometer2.6-1m in Arabidopsis roots. Data from 10-day-old seedlings expressing MGlucoMeter2.6-1m (Supplementary Video 1). Maximum intensity Z-projection and the emission ratio of cpsfGFP over LSSmApple are shown. Experiments were repeated three times independently with comparable results. Scale bar: 200 µm. Time-dependent *in vivo* response of MGlucometer2.6-1m in Arabidopsis roots.

### Detection of glucose accumulation by sucrose treatment to leaf

Sucrose generated by photosynthesis is loaded into the phloem of leaves and translocated from shoot to root. Sucrose utilization requires metabolic activities, e.g. invertases, that produce glucose and fructose. To investigate whether the MGlucoMeter2.6 can detect the production of glucose in roots that is derived from long distance translocation of sucrose from the shoots, shoots of Arabidopsis plants expressing MGlucoMeter2.6-1m were exposed to media containing sucrose, and glucose accumulation was monitored in root tips (Fig. S2). About 40 minutes after sucrose supply to the shoots, a ratio change was detected in roots, indicating that sucrose had been translocated from the shoot to the root tip unloading zone, where sucrose was then converted to glucose (Fig. 6, Supplementary Video 2). As one may expect, we did not observe a change in glucose levels in the vasculature, consistent with delivery of sucrose from the shoot in the phloem, followed by unloading in the unloading zone in the root and hydrolysis and diffusion in the unloading zone. Of note, MGlucoMeter2.6-1m did not show detectable responses even to high sucrose concentrations (Fig. 4). These observations are consistent with the important role of cytosolic invertases CINV1 and 2 for root growth and development (Barratt *et al*., 2009). In particular, CINV1 mRNA levels were high in cells in the unloading zone (Fig. S3). Thus, detection of glucose accumulation by sucrose treatment to shoot confirmed that the sensitivity (ΔR/R_0_ 180-230%) of MGlucoMeter2.6-1m is high enough for effective and sensitive wide-view imaging to explore long-distance sugar translocation and metabolism in Arabidopsis.

**Figure 6:**
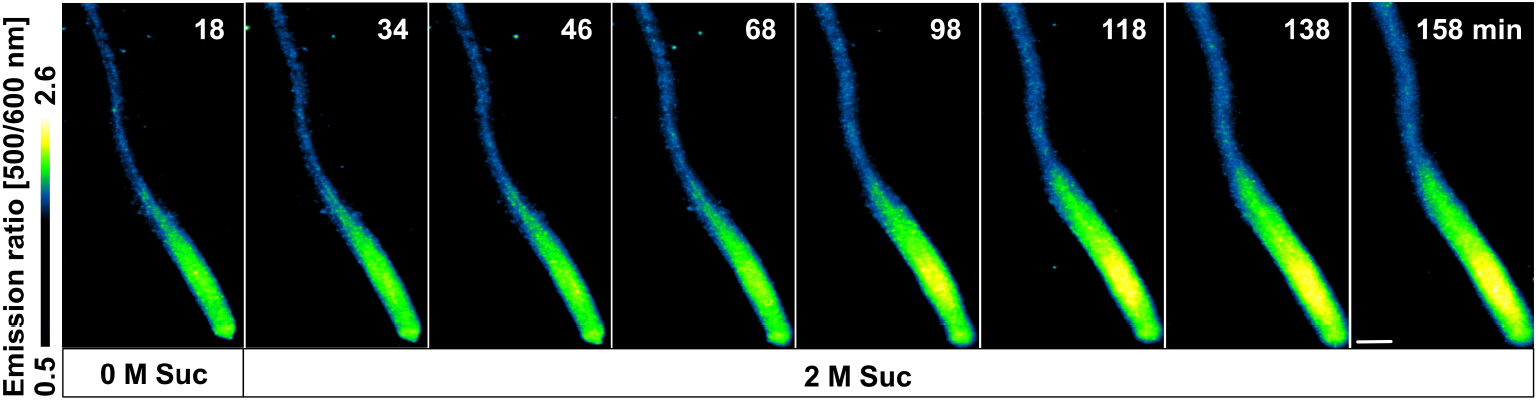
Detection of glucose accumulation in root tips of MGlucoMeter2.6-1m expressing Arabidopsis seedlings in response to sucrose application to the cotyledons. Time-dependent *in vivo* glucose accumulation in Arabidopsis roots after sucrose treatment to the cotyledons (Fig. S2). Data from 12-day-old seedlings expressing MGlucoMeter2.6-1m (Supplementary Video 2). Maximum intensity Z-projection images and the emission ratio of cpsfGFP over LSSmApple are shown. Scale bars: 200 µm. The experiment was repeated with three biological replicates (individual roots) and three times independently with comparable results.

In summary, we developed a new series of genetically encoded ultrasensitive ratiometric glucose sensors using the Matryoshka concept, in which excitation at a single wavelength enables detection of the emission for the sensory domain, a circularly permutated superfolder-GFP and the large stokes shift reference fluorophore LSSmApple. The sensory domain was a periplasmic glucose binding protein from a thermophile, which is likely more robust to insertion and mutagenesis. An alanine scan yielded sensors with a high sensitivity. We generated affinity mutants and used the optimized MGlucoMeter2.6 with an affinity for glucose of 1 mM to monitor glucose accumulation in intact Arabidopsis roots either from externally added glucose to roots or from hydrolysis of sucrose delivered from the shoot. Rapid hydrolysis of sucrose had previously been detected with a FRET glucose sensor (Chaudhuri *et al*., 2008).

Here we obtained evidence that sucrose is likely hydrolyzed after release from the phloem upon delivery from the shoot via the phloem. Cell-to-cell movement in the root is most likely mediated by plasmodesmata (Chaudhuri *et al*., 2008; Stadler *et al*., 2005; Wright and Oparka, 1997). The new sensors can likely be deployed in a wide range of organisms, as previously shown for the FRET glucose sensors, including bacteria, yeast and human cell lines (Bermejo *et al*., 2010; Bermejo *et al*., 2013; Bermejo, Haerizadeh, *et al*., 2011; Fehr *et al*., 2005; Takanaga *et al*., 2008).

## Author information

YI and WF conceived of the project. YI conducted all experiments. NZ and SP contributed to initial sensor construction. YI and WF prepared the figures. YI, NZ and WF wrote the manuscript. NZ and SP validated maps and sequences.

## Data availability

The raw data have been deposited as an ARC under doi (available for publication). Plasmids are available from Addgene (www.addgene.org).

## Acknowledgements

This work is part of the collaboration research center CRC1535 funded by grants to WF from Deutsche Forschungsgemeinschaft (DFG, German Research Foundation) - Project 458090666/CRC1535 and Germany′s Excellence Strategy – EXC-2048/1 – project ID 390686111 (CEPLAS), and an Alexander von Humboldt Professorship.

