## Supplementary material for "Evidence for rapid hydrolysis of shoot-derived sucrose using an ultrasensitive ratiometric Matryoshka-type MGlucoMeter sensor": Table S1

Table S1
Fig. S1
Fig. S2
Fig. S3
Supplementary Text
Supplementary Video 1
Supplementary Video 2

**Table S1. Primers for DNA construction and mutagenesis**

| Primer name | Sequence (5'→3') |
|---|---|
| Ishi797_MGlucoMeter2.0_f | Gttatattccagtttatgccccaga |
| Ishi799_MGlucoMeter2.0_r | aaactggaatataactttGCTaatcctaatgcatatggtcaaag |
| Ishi871_MGlucoMeter2.1_f | GCAggtcaaagtgcgatgcgcgactggcg |
| Ishi872_MGlucoMeter2.1_r | gcatcgcactttgaccTGCtgcattaggattgttaaagttatattccag |
| Ishi808_MGlucoMeter2.2_f | gctagcagggtatttggatgg |
| Ishi810_MGlucoMeter2.2_r | tccAAAtaccctgctGCACATaacgtgtatattaccgcggataaacagaa |
| Ishi806_MGlucoMeter2.3_f2 | cacgttTGCgctagcagggtatttggatggat |
| Ishi807_MGlucoMeter2.3_r2 | gctagcGCAaacgtgtatattaccgcggataaacaga |
| Ishi798_MGlucoMeter2.4_r | aaactggaatataacGCTAACaatcctaatgcatatggtcaaag |
| Ishi804_MGlucoMeter2.6_f | tgcatatggtcaaagtgcgatgcgcg |
| Ishi805_MGlucoMeter2.6_r | ctttgaccatatgcAGCaggattgttaaagttatattccagt |
| Ishi798_MGlucoMeter2.7_r | gatccatccAAAtacGCAGCtAGCCATaacgtgtatattaccgcggata |
| Ishi800_MGlucoMeter2.8_f | gttaaagttatattc |
| Ishi801_MGlucoMeter2.8_r | gaatataactttaacGCTcctaatgcatatggtcaaagtgcga |
| Ishi865_MGlucoMeter2.9_f | GCAcctgctagccataacgt |
| Ishi867_MGlucoMeter2.9_r | gttatggctagcaggTGCtttggatggatcgctatcaagacgtg |
| Ishi868_MGlucoMeter2.10_r | gctagcagggtaTGCggatggatcgctatcaagacgtgcag |
| Ishi802_MGlucoMeter2.12_f | accatatgcattAGCattgttaaagttatattccagttta |
| Ishi803_MGlucoMeter2.12_r | GCTaatgcatatggtcaaagtgcgatgcgcgac |
| Ishi811_MGlucoMeter2.13_r | gatccatccAAAtacGCAGCtAGCCATaacgtgtatattaccgcggata |
| Ishi804_MGlucoMeter2.15_r | GCTaatgcatatggtcaaagtgcgatgcgcgac |
| Ishi805_MGlucoMeter2.16_r | ctttgaccatatgcAGCaggattgttaaagttatattccagt |
| Ishi879_MGlucoMeter2.17_f | GCAcgcgactggcgcagtaaccggattgtagg |
| Ishi880_MGlucoMeter2.17_r | gcgccagtcgcgTGCcgcactttgaccatatgcattaggattgttaaagttatat |
| Ishi881_MGlucoMeter2.18_f | GCAagcgatccatccaaataccc |
| Ishi882_MGlucoMeter2.18_r | ggatggatcgctTGCaagacgtgcagcaatagagcctttcagtg |
| Ishi883_MGlucoMeter2.19_f | GCAgatccatccaaataccctgct |
| Ishi884_MGlucoMeter2.19_r | tttggatggatcTGCatcaagacgtgcagcaatagagcctttca |
| Ishi887_MGlucoMeter2.21_f | GCAtccaaataccctgctagccata |
| Ishi888_MGlucoMeter2.21_r | agggtatttggaTGCatcgctatcaagacgtgcagca |
| Ishi894_MGlucoMeter2.22_f | Gcgatgcgcgactggcg |
| Ishi1125_MGlucoMeter2.22_r | ccagtcgcgcatcgcTGCttgaccatatgcTGCagga |
| Ishi957_MGlucoMeter2.23_f | GCAaaataccctgctGCAcataacgtgtatattaccgcgg |
| Ishi890_MGlucoMeter2.23_r | agcagggtatttTGCtggatcgctatcaagacgtgcagc |
| Ishi892_MGlucoMeter2.25_f | ggtcaaagtgcgatgcgcg |
| Ishi952_MGlucoMeter2.25_r | catcgcactttgacctgctgcTGCaggattgttaaagttatat |
| Ishi953_MGlucoMeter2.26_r | cgcactttgTGCATAtgcTGCaggattgttaaagttatattccagt |
| Ishi875_MGlucoMeter2.27_f | GCAagtgcgatgcgcgactggcgcagtaac |
| Ishi954_MGlucoMeter2.27_r | gcgcatcgcactTGCaccatatgcTGCaggattgttaaagttata |
| Ishi955_MGlucoMeter2.28_r | gcgccagtcgcgTGCcgcactttgaccatatgcTGCaggattgttaaagttatat |
| Ishi956_MGlucoMeter2.29_r | ccatccaaataccctgctGCAcataacgtgtatattacc |
| Ishi889_MGlucoMeter2.30_f | GCAtccaaataccctgctagccata |
| Ishi924_H66A_f | tccataccggcCGCcacctgaaatgtgtccggaggatcaccgcc |
| Ishi1126_H66A_r | tgGCGgccggtatggaactcattggcacttgggtagttgcgaa |
| Ishi900_A66H_f | Atgccggtatggaactcattggcacttg |
| Ishi925_A66H_r | Gttccataccggcatgcacctgaaatgtgtccggaggatcaccg |
| Ishi898_H348A_f | gGCGggcgctgtcgctccggaaagtttcatg |
| Ishi899_H348A_r | gcgacagcgccCGCcactaagctgcctacaatccggttactg |
| Ishi1127_A348H_f | gCATggcgctgtcgctccggaaagtttcatg |
| Ishi1128_A348H_r | gcgacagcgccATGcactaagctgcctacaatccggttactg |
| Ishi1129_Inv_f | Ggcgctgtcgctccggaaagtttcatg |
| Ishi1130_MGlucoMeter2.6a_r | CGGAgcgacagcgccACGcactaagctgcctacaatccggttactg |
| Ishi1131_MGlucoMeter2.6-700μ_r | CGGAgcgacagcgccATTcactaagctgcctacaatccggttactg |

| | |
|---|---|
| Ishi1132_MGlucoMeter2.6b_r | CGGAgcgacagcgccATCcactaagctgcctacaatccggttactg |
| Ishi1133_MGlucoMeter2.6-7m_r | CGGAgcgacagcgccACAcactaagctgcctacaatccggttactg |
| Ishi1134_MGlucoMeter2.6c_r | CGGAgcgacagcgccTTGcactaagctgcctacaatccggttactg |
| Ishi1135_MGlucoMeter2.6d_r | CGGAgcgacagcgccTTCcactaagctgcctacaatccggttactg |
| Ishi1139_MGlucoMeter2.6e_r | CGGAgcgacagcgccCTTcactaagctgcctacaatccggttactg |
| Ishi1140_MGlucoMeter2.6f_r | CGGAgcgacagcgccAAAcactaagctgcctacaatccggttactg |
| Ishi1141_MGlucoMeter2.6g_r | CGGAgcgacagcgccAGGcactaagctgcctacaatccggttactg |
| Ishi1142_MGlucoMeter2.6h_r | CGGAgcgacagcgccACTcactaagctgcctacaatccggttactg |
| Ishi1143_MGlucoMeter2.6i_r | CGGAgcgacagcgccAGTcactaagctgcctacaatccggttactg |
| Ishi1144_MGlucoMeter2.6j_r | CGGAgcgacagcgccCCAcactaagctgcctacaatccggttactg |
| Ishi1145_MGlucoMeter2.6k_r | CGGAgcgacagcgccATAcactaagctgcctacaatccggttactg |

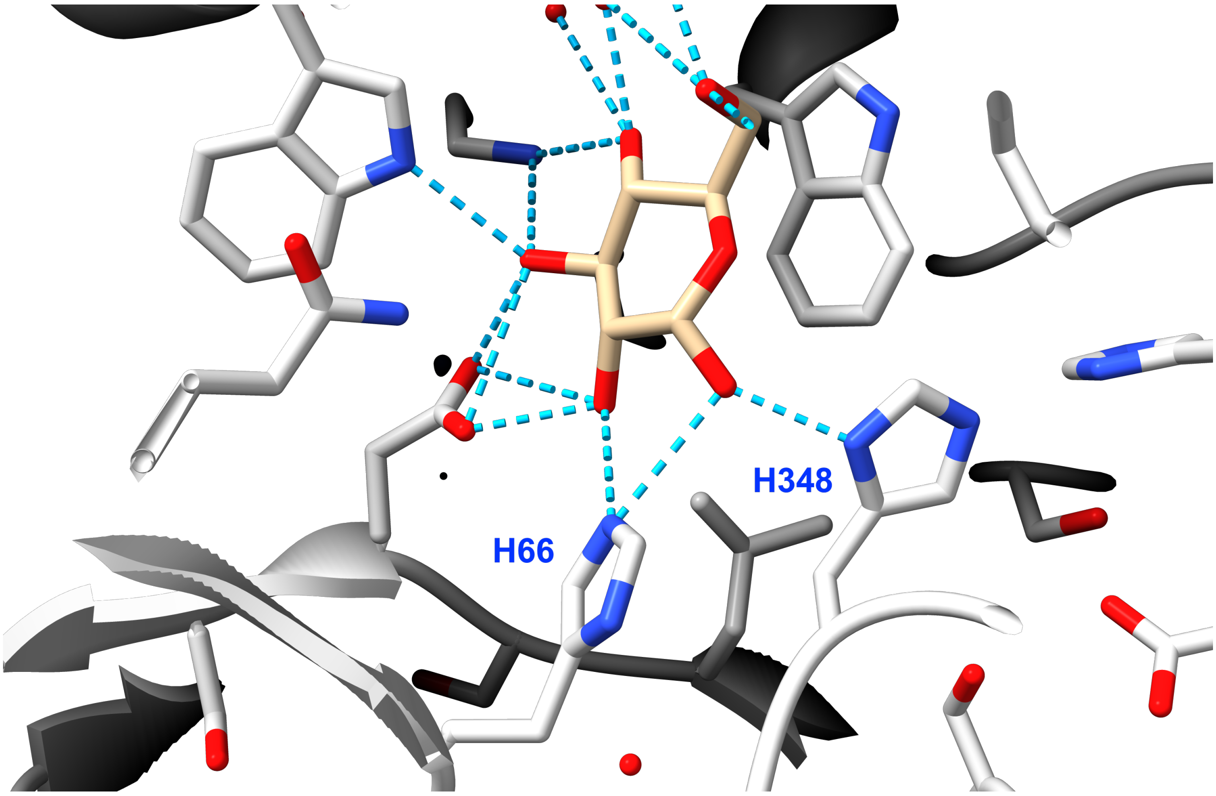


**Figure S1. Hydrogen bond interactions of two histidines with glucose in ttGBP (pdb 2B3B).** Hydrogen bonds between ttGBP and glucose are shown in turquoise. H66 and H348 are marked. The figure was generated using UCSF Chimera X (www.cgl.ucsf.edu/chimerax/).

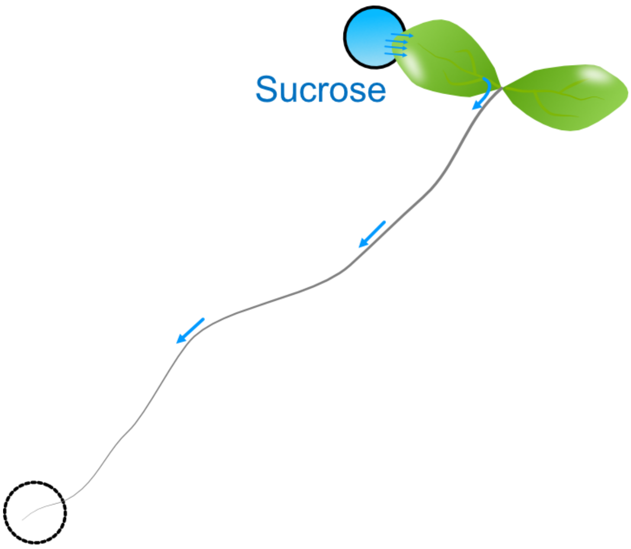


**Figure S2: Experimental setup for observing  glucose production in root tips.**

ROI used for quantification of the emission ratio in Fig. 6 is shown as dotted black circle. The tip of leaf was exposed to 2M sucrose containing ½ salt strength MS and 2.35 mM MES (pH 5.7) buffer. Roots were placed on 1% agar plates containing ½ salt strength MS and 2.35 mM MES (pH 5.7).

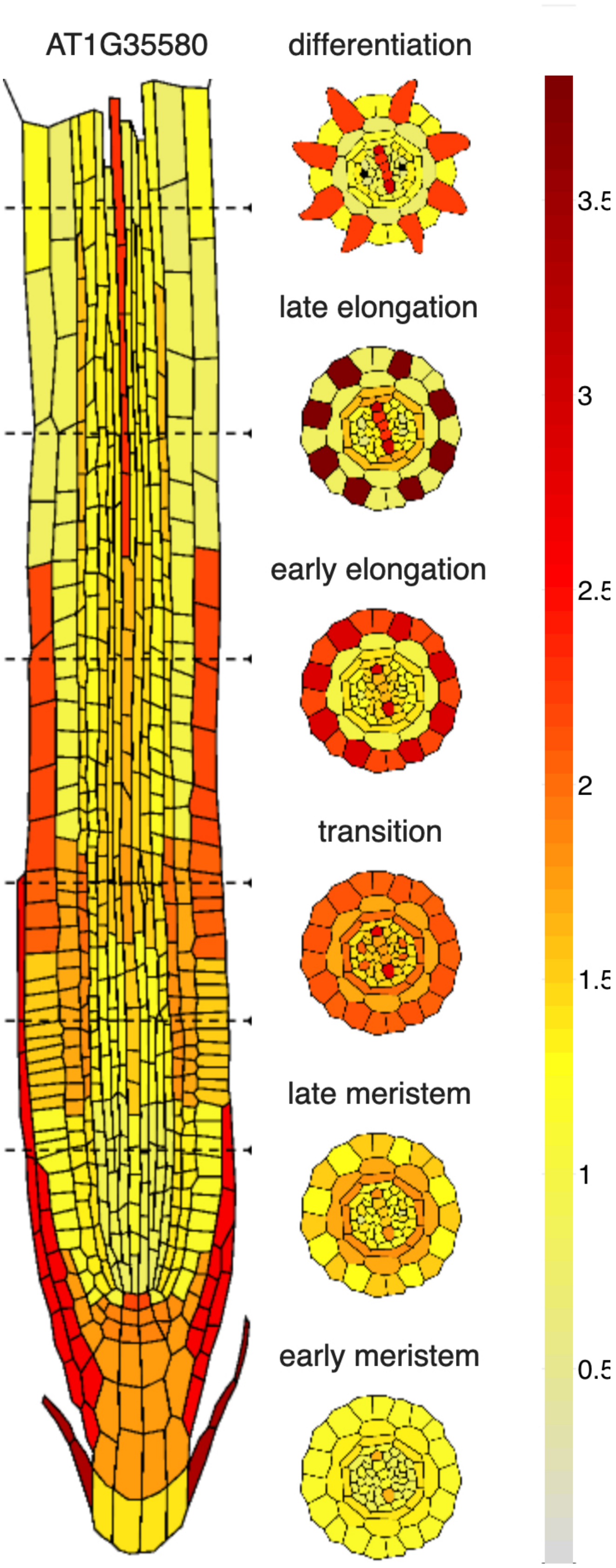


**Figure S3: Levels of cytosolic invertase CINV1 mRNA in cell types in the root tip derived from single cell sequencing data.**

Image downloaded from the *Root Cell Atlas* (https://rootcellatlas.org/). Generated by Rahul Shaw, Maria Savina, Xin Tian, Zhichao An, Victoria Mironova, and Jian Xu.

**Supplementary Text**

**Protein sequence of the ttgbp polypeptide from *Thermus thermophilus* used for sensor construction.**

The expressed gene product (P0328) lacks the 21 bp leader (Cuneo *et al.*, 2009)

Leader 1-21 MRKWLLAIGM VLGLSALAQG G

Protein used for structure determination (pdb: 2B3B)

```
MKLEIFSWWA GDEGPALEAL IRLYKQKYPG VEVINATVTG GAGVNARAVL  50
KTRMLGGDPP DTFQVHAGME LIGTWVVANR MEDLSALFRQ EGWLQAFPKG 100
LIDLISYKGG IWSVPVNIHR SNVMWYLPAK LKEWGVNPPR TWDEFLATCQ 150
TLKQKGLEAP LALGEN WTQ QHLWESVALA VLGPDDWNNL WNGKLKFTDP 200
KAVRAWEVFG RVLDCANKDA AGLSWQQAVD RVVQGKAAFN VMGDWAAGYM 250
TTTLKLKPGT DFAWAPSPGT QGVFMMLSDS FGLPKGAKNR QNAINWLRLV 300
GSKEGQDTFN PLKGSIAARL DSDPSKYNAY GQSAMRDWRS NRIVGSLVHG 350
AVAPESFMSQ FGTVMEIFLQ TRNPQAAANA AQAIADQVGL GRLGQ
```

Numbering of residues based on protein P0328
